# Machine learning-based spike sorting reveals how subneuronal concentrations of monomeric Tau cause a loss in excitatory postsynaptic currents in hippocampal neurons

**DOI:** 10.1101/2024.02.29.582792

**Authors:** Marius Brockhoff, Jakob Träuble, Sagnik Middya, Tanja Fuchsberger, Ana Fernandez-Villegas, Amberley Stephens, Miranda Robbins, Wenyue Dai, Belquis Haider, Sulay Vora, Nino F Läubli, Clemens F Kaminski, George G Malliaras, Ole Paulsen, Gabriele S Kaminski Schierle

**Affiliations:** Department of Chemical Engineering and Biotechnology, University of Cambridge; Department of Physiology, Development and Neuroscience, University of Cambridge; Electrical Engineering Division, Department of Engineering, University of Cambridge

## Abstract

Extracellular recordings of neuronal activity constitute a powerful tool for investigating the intricate dynamics of neural networks and the activity of individual neurons. Microelectrode arrays (MEAs) allow for recordings with a high electrode count, ranging from 10s to 1000s, generating extensive datasets of neuronal information. Furthermore, MEAs capture extracellular field potentials from cultured cells, resulting in highly complex neuronal signals that necessitate precise spike sorting for meaningful data extraction. Nevertheless, conventional spike sorting methods face limitations in recognising diverse spike shapes, thereby constraining the full utilisation of the rich dataset acquired from MEA recordings. To overcome these limitations, we have developed a machine learning algorithm, named *PseudoSort*, which employs advanced self-supervised learning techniques, a distinctive density-based pseudo-labelling strategy, and an iterative fine-tuning process to enhance spike sorting accuracy. Through extensive benchmarking on large-scale simulated datasets, we demonstrate the superior performance of *PseudoSort* compared to recently developed machine learning-based (ML) spike sorting algorithms. We showcase the practical application of *PseudoSort* by utilising MEA recordings from hippocampal neurons exposed to subneuronal concentrations of monomeric Tau, a protein associated with Alzheimer’s disease (AD). Our results, validated against patch clamp experiments, unveil that monomeric Tau at subneuronal concentrations induces stimulation-dependent disruptions in both local and global activity of hippocampal neurons. Remarkably, patch clamp electrophysiology highlights the effect of combined Tau and neuronal stimulation treatment on excitatory postsynaptic currents, whereas *PseudoSort* excels in identifying neuronal clusters that exhibit diminished firing capacity following Tau treatment alone, *i*.*e*., in the absence of stimulation. This comprehensive approach validates the prowess of *PseudoSort* and unravels the intricate effects of Tau on neuronal activity, particularly in the context of AD.

## Introduction

Microelectrode arrays (MEAs) have revolutionised the landscape of neuroscience research by enabling prolonged and extensive monitoring of local field potentials from neurons over a large area. This technology provides non-invasive recordings, capturing diverse spatial and temporal neuronal signals, thus providing additional information as compared to other electrophysiological techniques, such as patch-clamping (Obien *et al*., 2015). Despite these advantages, MEAs encounter challenges in both fundamental and therapeutic neuroscience research (Buzsáki, Anastassiou and Koch, 2012; Spira and Hai, 2020). These challenges arise from the inherent complexity of neuronal signal analysis, given that each electrode records signals from multiple neurons and various noise sources (Brown, Kass and Mitra, 2004; Buzsáki, Anastassiou and Koch, 2012; Quiroga, 2012; Anastassiou, Buzsáki and Koch, 2013), resulting complicated datasets.

A critical bottleneck in the analysis of MEA-recorded data is spike sorting, the process of attributing recorded spike signals, *i*.*e*., extracellular action potentials, to individual source neurons (typically up to six or seven per electrode (Quiroga, 2012)). This step is pivotal for enhancing our understanding of neuronal function and dysfunction (Quiroga, 2012; Rey, Pedreira and Quian Quiroga, 2015; Carlson and Carin, 2019). Despite the existence of a variety of spike sorting methodologies (Harris *et al*., 2000; Quiroga, Nadasdy and Ben-Shaul, 2004; Rutishauser, Schuman and Mamelak, 2006; Kadir, Goodman and Harris, 2014; Rossant *et al*., 2016; Chung *et al*., 2017; Yger *et al*., 2018; Buccino *et al*., 2020; Pachitariu, Sridhar and Stringer, 2023), limitations for example in accuracy, reliability, scalability and reproducibility (Gibson, Judy and Markovic, 2012; Rey, Pedreira and Quian Quiroga, 2015; Buccino, Garcia and Yger, 2022) still persist. Accurately identifying the number of present neurons and distinguishing their spikes becomes particularly challenging in densely packed neuronal cultures or brain tissue, where signal overlap is common or when neurons fire so rarely that they get lost between more abundant signals (Shoham, O’Connor and Segev, 2006; Quiroga, 2012). Moreover, the intrinsic variability in spike shapes, electrode drift or damage, and the presence of noise further complicate accurate predictions (Quiroga, Nadasdy and Ben-Shaul, 2004; Buccino, Garcia and Yger, 2022). Hence, addressing these challenges is crucial for advancing the capabilities of MEAs and unlocking their full potential in unravelling the intricacies of neuronal activity.

Machine learning (ML) approaches, with their capacity to handle large datasets and learn complex patterns, are well-suited to address the complexity of spike sorting. Recently, such data-driven ML approaches to spike sorting (Wu *et al*., 2018, 2019; Carlson and Carin, 2019; Lee *et al*., 2020; Li *et al*., 2020; Rácz *et al*., 2020; Eom *et al*., 2021; Rokai *et al*., 2021; Toosi, Akhaee and Dehaqani, 2021; Valencia and Alimohammad, 2021; Wouters, Kloosterman and Bertrand, 2021; Buccino, Garcia and Yger, 2022; Saif-ur-Rehman *et al*., 2023; Lu *et al*., 2024) have demonstrated for example improved accuracy and fast online processing. Here, we present a ML-based approach to spike sorting, *PseudoSort*, proposing a paradigm shift that includes self-supervised learning, data augmentation, and a pivotal density-based pseudo-labelling strategy. The method is benchmarked on large-scale simulated (Camuñas-Mesa and Quiroga, 2013; Lu *et al*., 2024) datasets, demonstrating superior performance in comparison to existing spike sorting algorithms.

We demonstrate the capabilities of *PseudoSort* in an *in vitro* neuronal culture model of neurodegeneration. In this model, primary hippocampal neurons are exposed to monomeric Tau, a protein intricately associated with Alzheimer’s disease (AD) (Mandelkow and Mandelkow, 1998), using MEAs. Notably, Tau, typically an intracellular protein, exhibits prion-like transfer between cells (Kfoury *et al*., 2012; Wu *et al*., 2016) as demonstrated in previous studies (Goedert *et al*., 2014; Sanders *et al*., 2014). Furthermore, we have previously established that monomeric extracellular Tau uptake alone is sufficient to induce Tau pathology (Michel *et al*., 2014). Hence, the pivotal question is whether such extracellular monomeric Tau is also capable of inducing neuronal signalling defects in primary hippocampal neurons. This prompted us to expose the cells to subneuronal concentrations of monomeric Tau, *i*.*e*., concentrations which are below the physiological concentration of Tau in neurons (~2 μM) (Butner and Kirschner, 1991; Khatoon, Grundke-Iqbal and Iqbal, 1992), and evaluate the outcomes using MEAs and *PseudoSort*. Additionally, analogous patch clamp experiments are performed to compare and confirm the MEA results.

Our findings demonstrate that 1 μM extracellular Tau induces activity-dependent disturbances in both the local and global activity of hippocampal neurons. Intriguingly, the patch clamp data unveil the effects of combined Tau and stimulation treatment on excitatory postsynaptic currents, while *PseudoSort* excels in detecting neuronal clusters that experience a loss of firing capacity following Tau treatment without further stimulation. Our integrated approach provides comprehensive insights into the nuanced impact of Tau on neuronal activity, offering valuable contributions to our understanding of AD-related mechanisms. Further, the significance of this research extends widely, providing a resilient analysis tool for neuroscientists across diverse research domains. Its applications span from fundamental inquiries into neural network dynamics to practical investigations in neurodegenerative disease models and the exploration of brain-computer interfaces. Finally, it sheds light on the intricate relationship between heightened neuronal activity and tauopathy, offering valuable insights into the underlying mechanisms of AD.

## Methods

### PseudoSort: Self-supervised density-based pseudo labelling for Spike Sorting

*PseudoSort* encompasses three main steps (**Fig. 1a**). First, an encoding model is pretrained in a self-supervised manner. Subsequently, preliminary labels, *i*.*e*., *pseudo labels*, are generated. Here, pseudo labels represent an estimated labelling that is built on sampling a high-confidence fraction of the dataset. Sampling such a fraction is based on the idea that some samples in the dataset which preserve a high local density, *i*.*e*., samples likely located at the centre of a cluster, are comparably easier to cluster than the full dataset. By sampling a comparably high-density fraction in a way that still preserves features representing the full dataset, pseudo-labelled samples are created which are subsequently used for fine-tuning.

**Figure 1:**
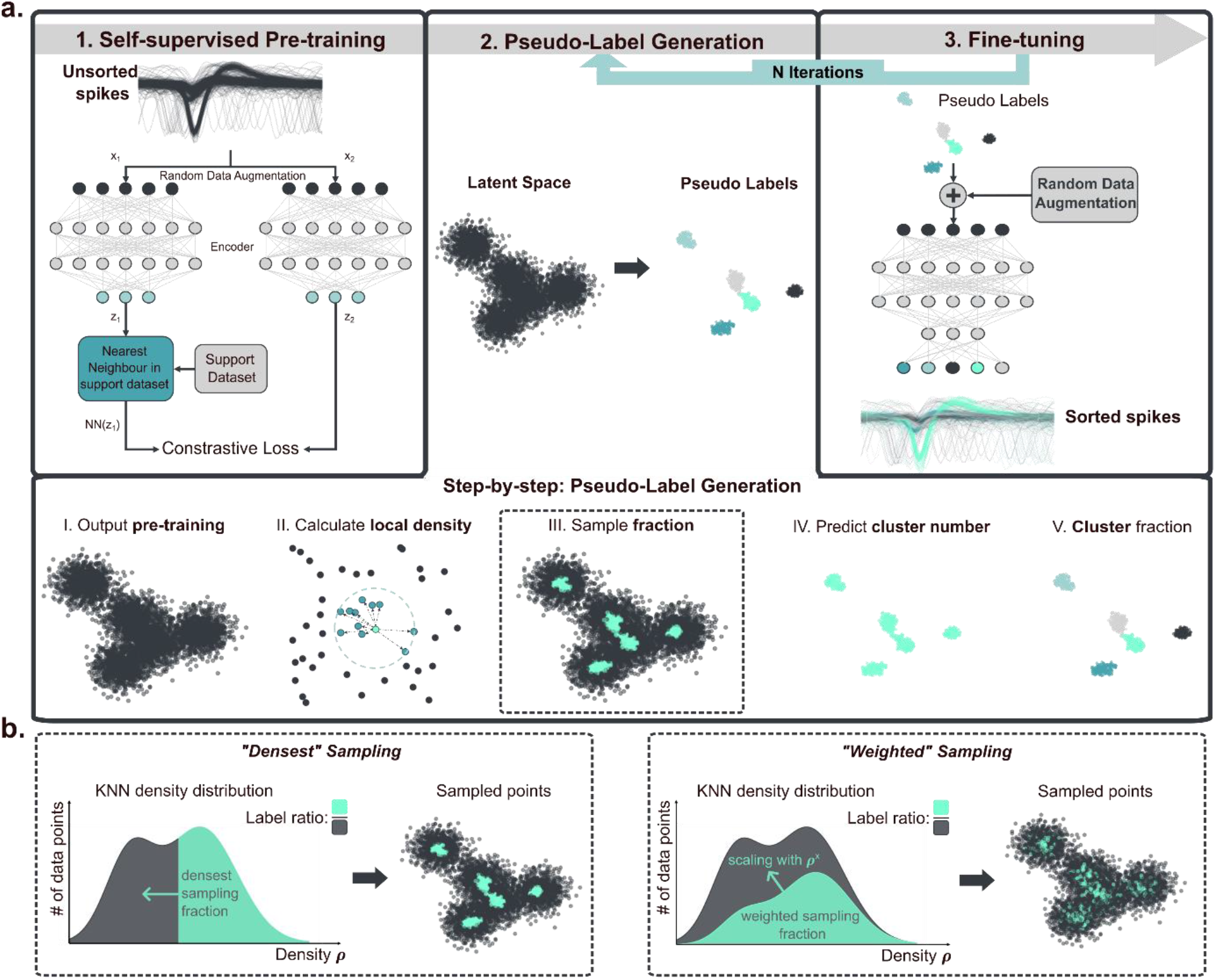
Schematic workflow of *PseudoSort*. **(a)** Top row: 3-step workflow of *PseudoSort*. Self-supervised pretraining (Dwibedi *et al*., 2021) on unsorted spike shape recordings yields an encoding model that produces a representative latent space. Based on the latent space, an iterative process of pseudo label generation and fine-tuning is executed, ultimately leading to high-accuracy classification for each spike shape recording. In the fine-tuning step, the encoding model is trained on the previously generated pseudo labels (semi-supervised problem) in a classification model. Lower row: detailed 5-step workflow to generate pseudo labels from latent space. Pseudo labels are sampled from whole latent space based on the local KNN density of each sample. The number of present clusters is predicted via the elbow method, and K-means++ is used to allocate all sampled points to one of the pseudo classes. **(b)** Two strategies to sample high-confidence samples from a dataset based on local KNN density. “Densest” describes the naïve approach of sampling the densest fraction of the dataset. Alternatively, the “weighted” sampling strategy that inherently favours points of higher local density but still captures a representative fraction of the dataset is proposed.

#### Pre-training

In the initial phase of the proposed methodology (**Fig. 1a**), self-supervised pre-training is employed utilising the Nearest-Neighbour Contrastive Learning of Visual Representation (NNCLR) (Dwibedi *et al*., 2021), adapted for the one-dimensional domain of neuronal signal processing. The input, pre-processed but unclassified spike waveforms, are augmented via the addition of Gaussian noise. This perturbation simulates variations in the recorded spikes, thereby generating multiple representations of the same neuronal event. The augmented signals are then passed through a fully connected encoder, which embeds the input data into a latent space. Within this latent space, a contrastive loss function is computed, using the nearest neighbour from a supportive dataset as a positive instance. The primary objective is to build up an encoder that acquires invariant features reflective of both the introduced stochastic noise and the intrinsic variability among the spikes. The pre-training stage aims to establish a basic understanding of the spikes within the latent space.

#### Pseudo-label generation

Following pre-training, the spikes are represented in a latent space where similar spikes are expected to cluster together. For the generation of pseudo labels, a subset of points believed to possess well-defined embeddings within the latent space is sampled – these representative points serve as a basis to fine-tune the encoding. Instead of clustering the full dataset at once, the aim is to identify a subset of the data that is inherently easier to classify.

The generation of pseudo labels involves several sub-steps (**Fig. 1a, lower row**). First, the local density of the latent space is calculated via the inverse of the average distance to K nearest neighbours (KNN density), setting K to 0.5% of the dataset size. Given a sampling fraction of the pseudo labels, two different sampling methods can be used (**Fig. 1b**). “Densest” sampling describes the naïve approach of sampling the densest fraction of the dataset. This strategy has the disadvantage that the densest fraction of the dataset is less likely to represent the full dataset and, therefore, might generalise poorly in the fine-tuning step. Instead, a “weighted” sampling method is proposed. Here, an exponential decay is overlayed with the normed density distribution. This leads to a high relative sampling chance for high-density points, while maintaining a non-zero sampling chance for even the least dense points. Biasing towards high-density points, *i*.*e*., high confidence as assumed in the centre of the clusters, ensures that the subsequent pseudo labels reflect the intrinsic distribution of the data, encompassing both high-density regions corresponding to frequent spike types and lower-density areas where rarer or unique spike patterns may reside. Using K-means++ (Arthur and Vassilvitskii, 2007) clustering on the sampled points, pseudo labels are assigned, *i*.*e*., a preliminary neuron class is assigned to each point within the sub latent space. These pseudo labels serve as proxies for the true classes, effectively transforming the unsupervised learning problem into a semi-supervised one. As K-means++ necessitates the number of distinct clusters, the elbow method (Thorndike, 1953; Syakur *et al*., 2018) is utilised to estimate the number of source neurons on 50% of the dataset, selected by weighted sampling.

Sampling a certain fraction of the dataset for high-confidence pseudo labels poses the challenge of identifying a fraction that works well for every dataset. It is demonstrated that choosing one fixed fractional value is not a feasible approach as different datasets exhibit different fractional pseudo label qualities (**Fig. 2a**) for either sampling method. To solve this problem, an iterative process is introduced, alternating between the steps of pseudo label generation and fine-tuning with an increasing fraction of sampled pseudo labels. As shown in **Fig. 2b**, this iterative approach achieves high-accuracy results for diverse datasets. Even if high accuracy is already achieved early (**Fig. 2b Small_1, Small_8**), this accuracy can be maintained and is not lost with further fine-tuning. At the same time, other datasets require the full range of steps and incrementally increase the overall performance over time (**Fig. 2b *Complex_12_1, Complex_14_2***). In this way, improved accuracy compared to choosing a fixed fractional value is achieved while reducing the required parameter choice.

**Figure 2:**
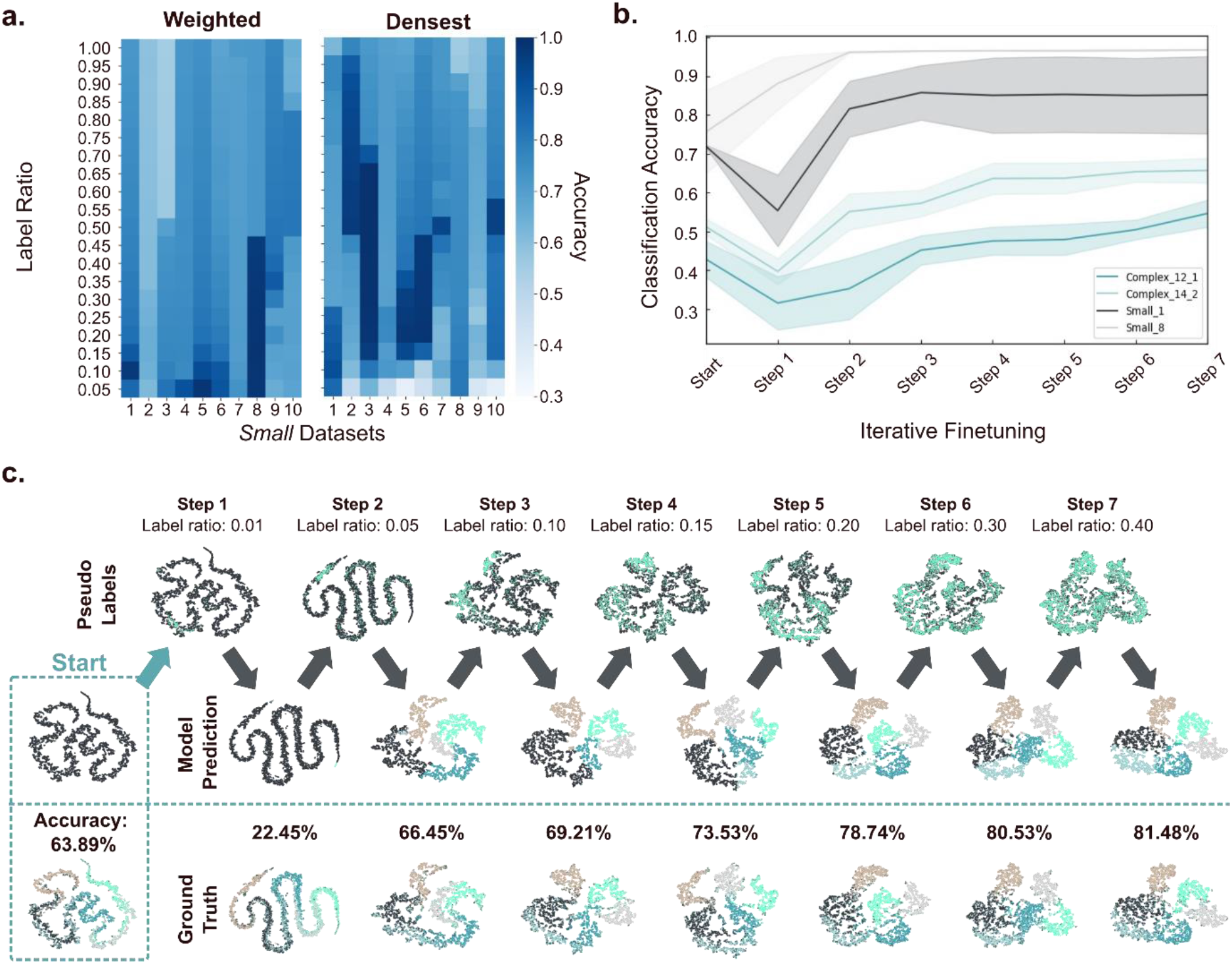
Pseudo label generation and fine-tuning iteratively improve spike sorting accuracy. **(a)** Heatmap showing the classification accuracy of sampled points for pseudo label generation for the *Small* datasets of “weighted” (left) and “densest” (right) sampling method. The y-axis describes the fraction of the dataset that is sampled, and accuracy describes the measured K-means++ classification accuracy achieved on the sampled data for each sampling method. **(b)** Measured accuracy during iterative pseudo label generation and fine-tuning for different example datasets (Complex_12_1, Complex_14_2, Small_1 and Small_8). Shown are mean (dark lines) and standard deviation (shaded area) for five repeated runs per dataset. **(c)** Illustrations of the iterative fine-tuning process with seven steps showing incremental improvements in model accuracy and the respective two-dimensional tSNE (Maaten and Hinton, 2008) visualisation of the latent space. Start with unlabelled latent space produced by pretraining. Top row indicates the sampled subset of the dataset of pseudo labelling, middle row shows the predicted classes of the model at each step and lower row shows the corresponding ground-truth classes. The initial accuracy and stepwise accuracies are given for each step. Dataset: *Complex_6_3*.

#### Fine-tuning

For fine-tuning, a final classification layer representing the different neuron classes is attached to the pre-trained encoder. With the classification layer in place, the encoder network is fine-tuned on the sampled pseudo labels. Random data augmentation is reintroduced to ensure the robustness and generalisation of the model.

Once the initial fine-tuning is completed, the model’s latent space is updated, and a new set of pseudo labels is sampled. These labels are then used for subsequent rounds of fine-tuning. This iterative process is repeated seven times, with each iteration expanding the ratio of pseudo labels (increasing pseudo label ratios: 0.01, 0.05, 0.10, 0.15, 0.20, 0.30, 0.40). By progressively increasing the number of labels, the model incrementally refines its understanding of the data, enhancing the granularity and accuracy of the spike sorting. The full process of iterative pseudo label generation and subsequent fine-tuning is illustrated for an example dataset (*Complex_6_3*) in **Fig. 2c**.

## Results

### *PseudoSort* benchmarking study on simulated data shows improved spike sorting performance

We first investigate how scaling the amount of neural recording data, *i*.*e*., the number of spike shape recordings, by one order of magnitude, while maintaining the same complexity—in this case, featuring spikes from five neurons—affects spike sorting performance (**Fig. 3a)**. Shown are the classification accuracies across the *Small* and *Large* datasets comparing *PseudoSort* with recent, ML-based spike sorting algorithms: AE-Ensemble (Eom *et al*., 2021), ROSS (Toosi, Akhaee and Dehaqani, 2021), and IDEC (Guo *et al*., 2017; Lu *et al*., 2024). These methods were chosen for benchmarking as they employ a comparable input to output structure as *PseudoSort* and allow direct classification of isolated spike shape recordings. Each method was evaluated five times on each dataset, and the mean accuracy was calculated to ensure robust and reliable results. For *Small* datasets, the median accuracy of *PseudoSort* is 75.80%, slightly above IDEC’s 73.82% and AE-Ensemble’s 72.97%, and significantly above the ROSS method 54.93%. Applied to the *Large* datasets, *PseudoSort’s* median accuracy is substantially higher at 92.05%, compared to IDEC’s 85.12% and well above that of AE-Ensemble at 71.91% and ROSS, which posts a median accuracy of 56.26%. This outcome suggests that *PseudoSort* scales effectively with increased data volume, capitalising on the larger dataset to enhance classification precision.

**Figure 3:**
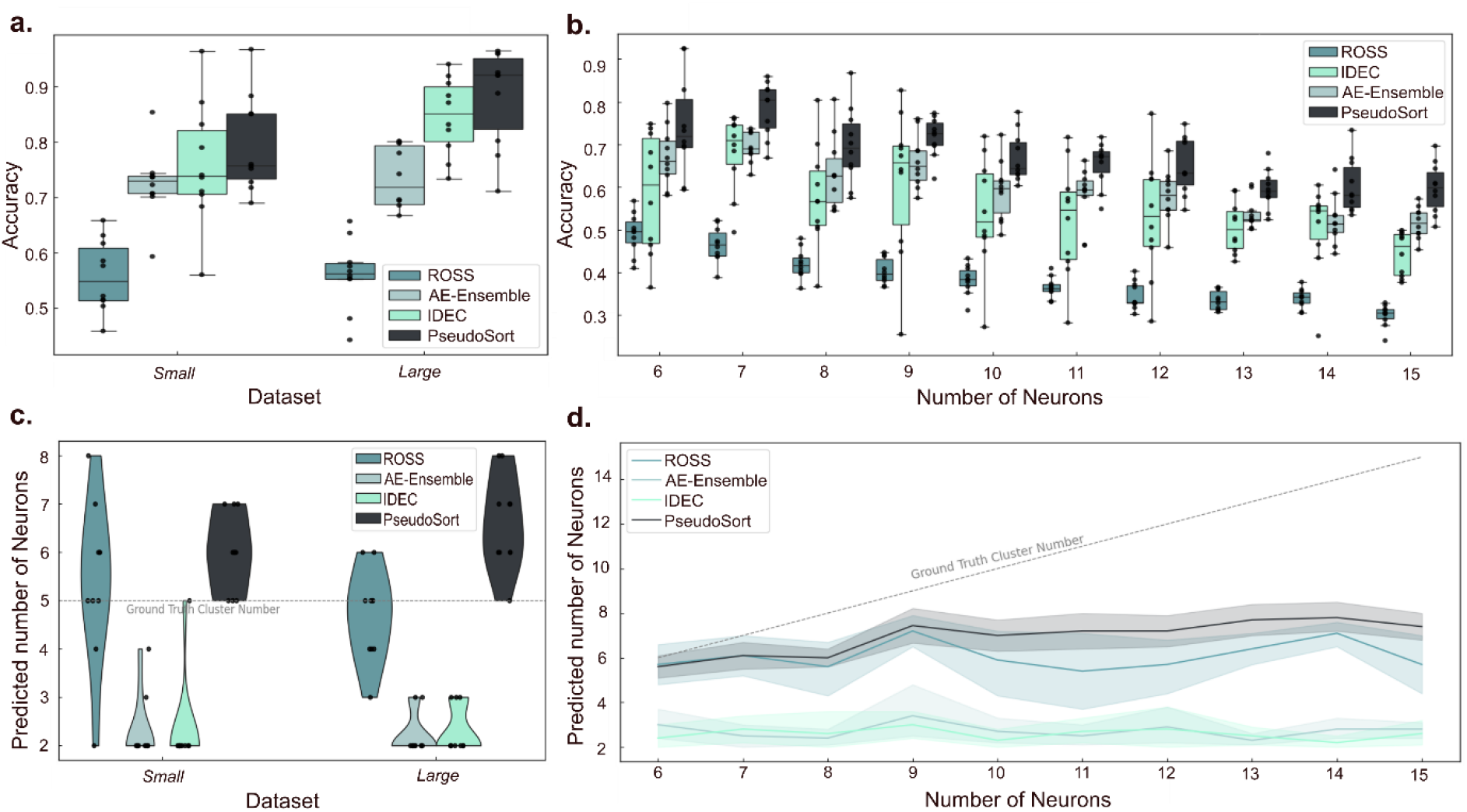
Evaluation of Spike Sorting accuracy and neuron number prediction across diverse data size and complexity levels illustrates *PseudoSort*’s superior performance. **(a)** Boxplots depicting the accuracy of spike sorting for *Small* and *Large* datasets, comparing *PseudoSort* with the AE-Ensemble (Eom *et al*., 2021), ROSS (Toosi, Akhaee and Dehaqani, 2021), and IDEC (Guo *et al*., 2017; Lu *et al*., 2024) methods. **(b)** Boxplots showing accuracy as a function of increasing number of neurons (*Complex* datasets), maintaining constant dataset size. **(c)** Violin plots representing the distribution of predicted neuron numbers for *Small* and *Large* datasets. **(d)** Trend lines with shaded interquartile range (IQR) areas illustrating the median predicted neuron numbers for all *Complex* datasets with the respective increasing number of neurons.

**Fig. 3b** explores the spike sorting performance when the complexity of the dataset is scaled up by increasing the number of neurons from 6 to 15 while keeping the size of the dataset constant. With the complexity scaled up to 7 neurons, *PseudoSort* outperforms the others, achieving a median accuracy of 80.39% compared to AE-Ensemble’s 68.98%, IDEC’s 70.94%, and ROSS’s 46.46%. This performance gap narrows as the complexity continues to increase, however, *PseudoSort* consistently leads in terms of median accuracy. For instance, with 10 neurons, *PseudoSort* registers a median accuracy of 64.64%, whereas the AE-Ensemble, IDEC, and ROSS methods drop to 59.63%, 51.93%, and 38.30%, respectively. Interestingly, at the highest complexity level with 15 neurons *PseudoSort* still preserves a relatively high median accuracy of 59.91%, whereas the AE-Ensemble and IDEC show only a slight decrease to 51.61% and 46.12% respectively, and ROSS declines more substantially to 30.50%. The trend across these varied complexity levels indicates that *PseudoSort* not only scales well with increased data volume, as shown **Fig. 3a**, but also demonstrates superior performance when faced with more intricate data structures, which is crucial for spike sorting applications.

The second important metric for the quality of a spike sorting approach is its ability to identify the correct number of signal sources, *i*.*e*., neurons, as shown in **Fig. 3c**. This is a critical aspect of spike sorting, as accurate identification of neuron numbers is essential for subsequent analyses. Across the five evaluations per dataset, the median predicted neuron number is shown. Again, the effect of scaling datasets in size is explored first. For the *Small* datasets, with a constant number of five neurons, the neuron predictions generated by *PseudoSort* predominantly converge around six neurons, indicating a tendency for overestimation of neuron numbers in certain scenarios. In contrast, the AE-Ensemble and IDEC methods demonstrate a notable underestimation of neuron numbers, with most of its predictions gravitating towards two neurons. Regarding ROSS, the predictions exhibit a broader distribution. The most frequent outcomes align with the accurate neuron count or exceed it by one. When increasing the number of spike samples (*Large* datasets, five neurons), the slight overestimation of neuron number of *PseudoSort* and significant underestimation of AE-Ensemble and IDEC persist, while ROSS’ predictions show less varied behaviour compared to smaller sample sizes.

Finally, the predictive performance of each method regarding neuron numbers is investigated as the complexity of the datasets increases **(Fig. 3d)**. Across all levels of complexity, it is observed that all methods tend to underestimate the number of neurons significantly. As dataset complexity grows, *PseudoSort* and ROSS exhibit a mild upward trend in predicted neuron numbers, indicating some adaptability to complexity. Conversely, IDEC and the AE-Ensemble predictions remain consistently low, showing no significant increase with greater complexity. Among the methods, *PseudoSort’s* median predictions are the least underestimated, with IDEC and the AE-Ensemble showing the most significant underprediction, and ROSS falling in between but displaying considerable variance. In conclusion, clear deficits in all spike sorting algorithms’ ability to predict the number of neurons for increasing number of neurons (>7) have been identified. Meanwhile, it has been shown that *PseudoSort* outperforms the existing alternatives in accurately clustering spike shape recordings as well as predicting the number of source neurons.

### Electrophysiological experiments unveil neuronal activity-dependent effects of monomeric Tau treatment at subneuronal concentrations

To assess the capability of *PseudoSort* in detecting neuronal defects, we subjected primary hippocampal neurons to subneuronal concentrations of monomeric Tau and monitored their activity using MEAs in the presence and absence of electrical stimulations. While Tau treatment alone, given the low concentration, did not lead to a measurable change in spike rate (**Fig. 4a**), a significant increase in spike rate was observed for cells that were electrically stimulated (**Supplementary Fig. 1**). However, intriguingly, in the presence of Tau, electrical stimulation failed to induce a spike rate increase as observed in the stimulation-only condition (**Fig. 4a**). To validate these findings, patch clamp recordings were performed using ChR2-YFP hippocampal neurons. Excitatory neurons (CA3 hippocampus) were optogenetically stimulated and, consecutively, their excitatory postsynaptic currents (EPSCs) were measured at EYFP-negative neurons. **Fig. 4b** illustrates a significant reduction in current during Tau + light stimulation compared to the light stimulation treatment, accompanied by distinct changes in the current profile at the respective neuron. And similar to the observations made from the MEA data, Tau-only treatment was not significantly different to the control without Tau, while light stimulation amplified the amplitude (**Supplementary Fig. 1**) and duration of the current profile.

**Figure 4:**
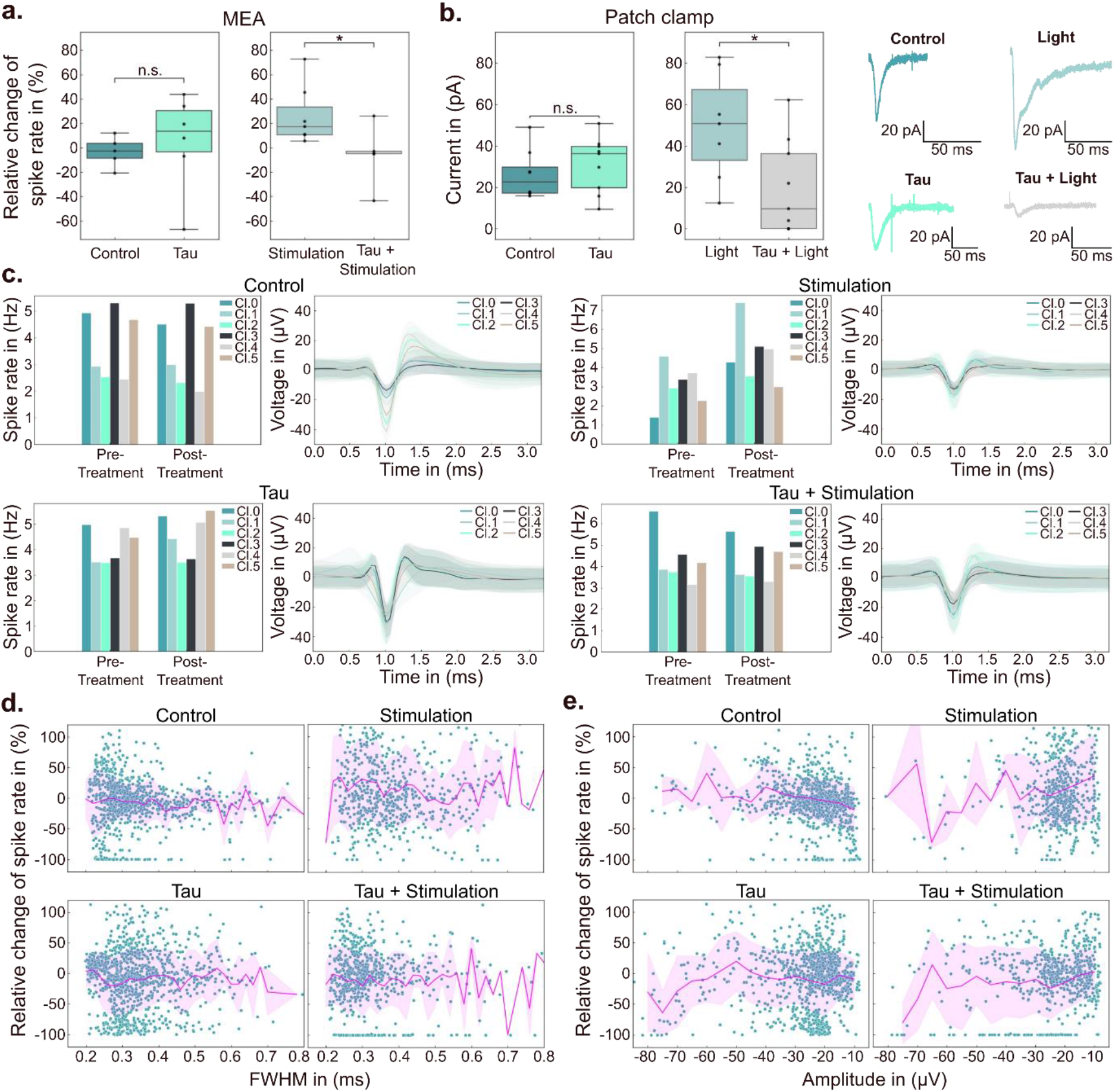
Electrophysiological experiments reveal neuronal activity-dependent effects of Tau. **(a)** Relative change of spike rate measured on MEAs following different treatments using Tau and electrical stimulation, compared to the pre-treatment baseline. N>=5. n.s. (not significant) P > 0.05; *P ≤ 0.05. **(b)** Patch clamp data of hippocampal neurons in the presence or absence of Tau treatment and light stimulated activity (left), and example traces showing EPSCs recorded from neurons after the different treatment/stimulation conditions (right). N>=7. n.s. (not significant) P > 0.05; *P ≤ 0.05. **(c)** Demonstration of single electrode analysis enabled by *PseudoSort*. Shown are sorted spike shape classes (Cl.0 – Cl.5, mean ± standard deviation as shaded area) found at single example electrodes and their respective firing rates before and after treatment for all conditions. **(d + e)** Explorative differential analysis enabled by spike sorting. Shown are relative changes of firing rate for every single spike class identified by *PseudoSort* as a function of the spike classes’ respective full width half maximum (FWHM) **(d)** or amplitude **(e)**. Each dot represents one specific spike class found at one electrode with mean and standard deviation shown as a magenta line and shaded area, respectively.

Insights from the patch clamp experiments are limited to single neurons. On the contrary, *PseudoSort* can provide further understanding at different scales (network vs. single neurons) by analysing spikes recorded at each electrode. **Fig. 4c** showcases examples where *PseudoSort* classified all spikes at each electrode in every experiment. The depicted example electrodes display typical spike shapes observed during the experiment, along with their respective spike rates before and after treatment. For these example electrodes, Tau + stimulation treatment induced only slight changes in spike rates for all classes, similar to Tau-only and the control. As anticipated, stimulation-only showed a general increase in spike rates for all spike classes at this specific electrode.

While analysing every single electrode provided insights into local effects, interpretation is challenging due to significant variations between electrodes. Therefore, **Fig. 4d** and **Fig. 4e** explore the possibility of aggregating single neuron effects to extrapolate overall trends. **Fig. 4d** depicts the relative change in spike rates for each single neuron class as a function of the average FWHM of the respective spike class, revealing width-dependent effects for Tau-only treatment. A drop to negative spike rates is observed for spikes with FWHM around 0.27 ms, indicating that many neurons largely or completely lost their activity in that regime. Similarly, **Fig. 4e** visualises the relative change in spike rates for each single neuron class as a function of the average amplitude of the respective spike class. A slight increase in spike rate after treatment is observed for spike classes with an amplitude of approximately −20 μV compared to smaller amplitude spikes (around −10 μV) for stimulation only, Tau-only, and Tau + stimulation, which is not observed for the no Tau control. Consistent with **Fig. 4d**, many neurons of amplitudes between −30 μV and −20 μV are largely lost (< −50% change in spike rate) for Tau-only treatment.

## Discussion

*PseudoSort* marks a significant shift in ML-based spike sorting methodologies by adopting self-supervised learning, aligning with recent advances in ML (Krishnan, Rajpurkar and Topol, 2022). This approach enhances the understanding of complex neuronal data, leading to more nuanced neuronal activity analysis. Its key feature, a density-based pseudo labelling strategy, ensures adaptability and generalisability by focusing on representative spike samples. Fine-tuning with pseudo labels crucially improves signal classification accuracy, especially for complex spike samples. An iterative fine-tuning process, designed to explore the space of pseudo labels progressively, avoids falling into local minima due to insufficient or poor-quality pseudo labels. The improvements from each iteration are demonstrable, with clearer clustering in the latent space and enhanced overall accuracies. Outperforming current ML-based spike sorting methods on simulated, single channel spike recording data, *PseudoSort* exhibits higher classification accuracy. This is particularly notable when scaling with increased data volumes or complexity, both of which are an important aspect given the capabilities of modern MEAs to record a vast amount of intricate neuronal signals (Steinmetz *et al*., 2021).

However, common challenges in the field of spike sorting are limited accuracy, especially for large number of neurons, remain (Buccino, Garcia and Yger, 2022). Here, the difficulty spike sorting algorithms face in accurately discerning neuron numbers in data-rich environments are demonstrated, with our method displaying marginal superiority under these challenging conditions. Although *PseudoSort’s* approach for detecting the number of neurons surpasses current alternatives, it is still limited, particularly when dealing with larger neuron populations. Anatomical considerations suggest that the number of neurons per electrode should be significantly higher (10x) than the generally detected up to six or seven units per electrode (Henze *et al*., 2000; Quiroga, 2012). The observed phenomenon could potentially stem from several factors, such as the prevalence of silent neurons induced by electrode-related tissue damage, or limitations inherent in current spike sorting algorithms, which may struggle to distinguish the activity of numerous neurons. Our analysis indicated that despite employing various spike sorting algorithms, including *PseudoSort*, none consistently identified more than eight source neurons. This observation suggests that the performance of spike sorting algorithms might serve as a significant bottleneck in detecting all neurons per electrode, underscoring the need for ongoing research in this area. It is imperative for new spike sorting methods to be extensively benchmarked, as satisfactory performance on small and simple datasets does not necessarily translate to more complex data, as obtained from biological measurements. It is foreseen that the datasets provided here can be utilised for future benchmarking studies and should undergo gradual expansion. The interpretation of spike sorting results can be challenging due to the intricate nature of neuronal data and the sophisticated algorithms employed. To address this issue, an easy-to-use script is provided, however, a fundamental understanding of the algorithms remains crucial for effective optimisation and troubleshooting. Moreover, reliance on simulated datasets for benchmarking algorithms may introduce biases, as these datasets may not comprehensively encompass the diversity of real neuronal data (Buccino, Garcia and Yger, 2022; Pachitariu, Sridhar and Stringer, 2023). Therefore, validation using a variety of experimental data and recognition of the limitations associated with simulated datasets are essential.

Furthermore, we strategically designed an experiment to assess the algorithm’s efficacy in extracting neuronal signalling data from intricate and challenging biological datasets. In particular, we wanted to address the molecular mechanisms underlying subneuronal concentrations of extracellular monomeric Tau by employing MEA and patch clamp techniques. Thus far, there has been no demonstration that low concentrations of extracellular monomeric Tau induce notable defects in neuronal signalling of hippocampal neurons. The here presented experimental findings reveal a distinctive response: Tau treatment during heightened neuronal activity, even at below-physiological concentrations, seems to mitigate the enhancing effects of the stimulation (**Fig. 4a, Supplementary Figure 1**), while Tau-only treatments fail to exhibit a significant opposing impact on neuronal activity. These MEA-based results are corroborated by patch clamp experiments, where Tau in combination with stimulation leads to a reduction in excitatory postsynaptic currents (EPSCs) in ChR2-YFP excitatory hippocampal neurons compared to stimulation only (**Fig. 4b**). Further, similar to the observations made with MEAs, Tau-only treatments in patch clamping fail to exhibit a significant impact on EPSCs while stimulation only causes a significant increase in EPSCs (**Supplementary Figure 1**). This observation implies that Tau, in conjunction with stimulation, specifically influences excitatory synapses, a phenomenon previously suggested in clinical contexts (Chang *et al*., 2021; Ranasinghe *et al*., 2022).

Utilising *PseudoSort*, we explore additional avenues of analysis to maximise the information gleaned from MEA recordings. **Figs. 4c-e** present outcomes facilitated by spike sorting on individual electrodes, demonstrating the capabilities of this approach. Here, it is illustrated that not all spike classes uniformly increased their spike rate, suggesting the ability to observe differential effects for each putative neuron. Combining network-level insights with data from single electrodes enables researchers to achieve a more comprehensive understanding at various levels (network → electrode → neuron), which has previously been suggested for the study of axonal dysfunction in neurodegenerative diseases (Yuan et al., 2020). Notably, *PseudoSort* proficiently extracts a subset of neurons in the Tau-only group that exhibit a decrease in spike rates to negative values for spikes with a FWHM of approximately 0.27 ms (**Fig. 4c, d**). This observation underscores a specific effect of exogenous Tau, consistent with findings from the patch clamp data on excitatory neurons. In general, patch clamp experiments are much more laborious (lower through-put) and only capable of recording single neurons. However, the patch clamp experiments alone have not been able to pick up a specific Tau-only treatment effects, highlighting the potential of *PseudoSort* to provide a more detailed analysis of the recorded data at much lower experimental expense. Nevertheless, for the specific dataset under consideration, the overall surge in activity during the stimulation condition cannot be unequivocally linked to either the width or amplitude of the corresponding spike class. Instead, an overarching observation emerges, wherein each neuron, on average, manifests heightened firing—a phenomenon mitigated by the inclusion of Tau during stimulation. This nuanced analytical approach is inferred to contribute significantly to a more comprehensive understanding and holds the potential to yield novel insights in forthcoming experiments. An additional illustration of the potential of *PseudoSort* lies in its ability to evaluate varying levels of synchronicity, as depicted in **Supplementary Fig. 2**. In this context, diverse scales of synchronicity, ranging from global (pertaining to interactions between all electrodes) to local (involving interactions among single neuron classes at each electrode), are delineated. These scales can be systematically compared to enrich our comprehension of the underlying effects (Singer, 1999; Uhlhaas and Singer, 2010).

*PseudoSort* makes significant contributions to advancing neuroscience by providing deeper insights into the dynamics of neural networks. This capability is invaluable for studying neurodegenerative diseases and mental disorders, enhancing our understanding of their underlying mechanisms. Future research in the field of self-supervised spike sorting is poised to pursue two distinct technical trajectories. Firstly, there is a drive to enhance accuracy in handling extensive datasets, facilitated by advancements in high-density MEAs. This may involve integrating transformer models and autoregressive training methods, leveraging abundant neuronal data to achieve greater precision. Secondly, there is a need to integrate self-supervised learning frameworks into more compact models, enabling efficient real-time analysis of neuronal data. We anticipate that these advancements in spike sorting methodologies will not only deepen our understanding of neuronal dynamics and diseases (Franke *et al*., 2012) but also catalyse advancements in brain-computer interface development by refining the interpretation of neuronal signals (Todorova *et al*., 2014).

## Supplementary Materials & Methods

### PseudoSort

#### Code and data availability

The presented methodology has been implemented on Python 3 using the TensorFlow (TensorFlow Developers, 2023) library. All codes and datasets will be made available upon publication.

#### Benchmark datasets

The datasets introduced here are designed to serve as a comprehensive, ground-truth framework for testing and comparing emerging spike-sorting algorithms. They comprise simulated recordings of spike shapes generated using NeuroCube (Camuñas-Mesa and Quiroga, 2013) in standard configuration (a single electrode, 300,000 neurons/mm^3^, 7% active neuron ratio, exponential firing rate distribution, and a 20 KHz sampling rate). In each recording, five neurons were positioned around a single electrode. The distance of each neuron to the electrode was determined by random selection within a range of 0 to 1 (in increments of 0.01), and their firing rates were randomly chosen to between 15 and 35 Hz. We provide the created Cube files, raw recording files in .mat file format as well as isolated spike shape recording files, stored as Python pickle files. Here, the first column carries the ground-truth spike class for each spike (integer). The second column contains the spike time of the stimulated spike (in ms). Finally, columns 3 to 66 contain the respective spike shape (64 data points). The datasets are organised in size and complexity. The group of sets called *Small* and *Large* consist of ten sets of spike recordings of five classes (source neurons) each, and include about 100.000 and 1.100.000 spike shape recordings, respectively. The *Complex* set group consists of 100 sets, each containing 100,000 spike shape recordings sourced from 6 to 15 neurons (ten sets each). These sets are formed by merging the ten *Small* datasets, from which a specific number of source neurons and their respective recordings are randomly chosen. Subsequently, the spike shape recordings are randomly sampled to create these sets.

#### Model training

*PseudoSort*’s encoder is a fully connected neural network with a 10-dimensional latent bottleneck. The encoder consists of dense layers of dimension [63, 500, 500, 2000, 10] with ReLU activation functions. For the NNCLR pre-training phase, the autoencoder is trained for 25 epochs (with only 25% of the dataset, randomly chosen) using Adam optimizer (Kingma and Ba, 2014), learning rate of 1e-3, batch size of 256, temperature of 0.1, queue size of 0.1 (relative to the dataset size, i.e. 10%) and projection width of 10. For the contrastive augmenter, Gaussian noise with max noise level of 0.075 are used, whereas the classification augmenter uses a max noise level of 0.04. While multiple data augmentation approaches (Wen *et al*., 2021) have been tested, Gaussian noise produced the most reliable outputs. For fine-tuning, a batch size of 128 and 50 epochs are used per iteration and a max noise level of 0.1 for the classification augmenter. The pseudo labels are generated iteratively, sampling increasing pseudo label ratios (0.01, 0.05, 0.10, 0.15, 0.20, 0.30, 0.40) of the dataset and with 0.5% of dataset samples as k-nearest neighbours. All models were trained on a NVIDIA A-100-SXM-80GB GPU.

#### Source neuron prediction

In the case of all benchmarked methods, the range for identifying the optimal cluster count is limited from 2 to a maximum of 20 source neurons.

### Experiments stimulation-dependent Tau pathology

#### Dissection of primary rat neurons

Hippocampal neurons of Sprague-Dawley rats (Charles River, UK), at 2 days postnatal (P2), were excised and gathered into 2 ml Eppendorf tubes filled with cold DMEM (Sigma-Aldrich, UK), and kept chilled on ice. Following the collection of tissue, the initial cold DMEM was replaced with DMEM at room temperature, supplemented with 0.1% Trypsin and 0.05% DNase (Sigma–Aldrich, UK). The tubes were subsequently placed in a CO_2_ incubator maintained at 37°C, with 5% carbon dioxide and 20% humidity, for a duration of 20 minutes. The cells underwent four washes with DMEM containing 0.05% DNase at room temperature and were dispersed into a single-cell suspension through gentle pipetting, first with a 1 ml and then with a 200 μL Gilson pipette tip. Following centrifugation of the cell suspension at 600 rpm for 5 minutes, the supernatant was discarded, and the cell pellet was softly resuspended in DMEM containing 10% FBS. The number of cells was assessed using a haemocytometer.

#### Cell culture

After autoclaving, the MEAs were cleaned with ethanol and DI water (3x each). They were then incubated with poly-L-lysine (PLL) solution (Sigma-Aldrich, UK) overnight. On the next day, the PLL was rinsed off three times with Dulbecco’s Phosphate Buffered Saline (DPBS) and subsequently placed under an ultraviolet lamp in a sterile laminar flow cabinet for two hours in order to activate the surface. Then, 1mL of NbActiv4 growth medium (BrainBits, U.S.) was introduced. The devices were placed in an incubator set at 37°C, 5% carbon dioxide, and 20% relative humidity, to warm up before plating primary post-natal day 2 (P2) hippocampal neurons. 180,000 primary hippocampal cells were then plated directly onto the device. 200 *μ*L of media was taken out and replaced by 300 *μ*L of warmed up NbActiv4 medium every other day to maintain the cell culture in an incubator set at 37°C, 5% carbon dioxide, and 20% relative humidity until the devices were used for experiments on days *in vitro* (DIV) 21.

#### Expression and purification of htau40

The human microtubule-associated protein tau (htau40) was recombinantly expressed using the pET29b vector in the E. coli strain BL21(DE3)-CondonPlus-RIPL (Agilent, U.S.) after transformation. The plasmid was obtained from Addgene (Plasmid#16316). The protein expression was induced using IPTG (isopropyl β-d-1-thiogalactopyranoside) as previously described (Barghorn, Biernat and Mandelkow, 2004). Briefly, a single colony was inoculated into 5 mL of LB (Lysogeny Broth) with Kanamycin (50 μg/mL) and grown overnight at 37°C. Subsequently, 1 mL of the overnight culture was transferred to 400 mL of LB media and grown at 37°C with agitation at 220 rpm until an optical density at 600 nm (OD600) of 0.6 was reached, following which the culture was induced with 0.5 mM IPTG and further cultivated at 18°C for overnight. The cell pellet was collected by centrifugation at 10,000 g for 10 minutes, resuspended in MES buffer (20 mM MES (2-(N-morpholino) ethanesulfonic acid), 1 mM EGTA (ethylene glycol-bis(β-aminoethyl ether)-N,N,N′,N′-tetraacetic acid), 0.2 mM MgCl2 (magnesium chloride), 5 mM DTT (dithiothreitol), 0.1 mM PMSF (phenylmethylsulfonyl fluoride), pH 6.8) suppled with cOmplete^TM^ Protease Inhibitor Cocktail, and lysed using an Ultrasonic Processor XL sonicator (Heat Systems, U.S.) with a cycle of 5 s on and 5 s off for a total of 5 minutes until the suspension became less opaque. The cell lysate was supplemented with extra NaCl (sodium chloride) to a final concentration of 0.5 M and further boiled in a water bath at 90°C for 20 minutes before cooling on ice. Subsequently, the cell lysate was centrifuged at 30,000 g for 20 minutes before the supernatant was poured out and filtered through a 0.45 μm Sartorius Minisart NML syringe filter. The soluble fraction was dialysed overnight at 4°C in a dialysis buffer (20 mM MES, 1 mM EGTA, 1 mM MgCl2, 2 mM DTT, 0.1 mM PMSF, pH 6.8 using NaOH (sodium hydroxide)) as to make sure that the concentration of NaCl is less than 10 mM, before proceeding with the purification step.

To check the presence of htau40 in the soluble fraction after dialysis, the samples were combined with SDS loading buffer (20% glycerol, 100 mM Tris–HCl (tris(hydroxymethyl)aminomethane-hydrochloride), 4% SDS (sodium dodecyl sulfate), and 0.2% bromophenol blue, pH 6.8) and boiled at 90°C for 10 minutes and loaded onto NuPAGE® Novex® 10% Bis-Tris Protein Gels (ThermoFisher Scientific, UK) and run for 45 minutes at 180 V. The dialysed protein underwent purification via ion exchange chromatography (IEX) against a linear gradient of buffer B on a HiPrep Q FF 16/10 anion exchange column (GE Healthcare, Sweden) in buffer A. Buffer A consisted of 20 mM MES, 1 mM EGTA, 1 mM MgCl2, 2 mM DTT, 0.1 mM PMSF, pH 6.8, while buffer B contained 20 mM MES, 1 mM EGTA, 1 mM MgCl2, 2 mM DTT, 0.1 mM PMSF, 250 mM NaCl, pH 6.8. The elution fraction was further purified using size exclusion chromatography (SEC) on a Superdex 200 16/60 column in HEPES buffer (10 mM HEPES (4-(2-hydroxyethyl)-1-piperazineethanesulfonic acid), 100 mM NaCl, 1 mM TCEP (Tris(2-carboxyethyl)phosphine), pH 7.5). The purification process was carried out on an ÄKTA Pure (GE Healthcare, U.S.) and htau40 was concentrated using 10k MWCO Amicon centrifugal filtration devices (Merck KGaA, Germany). The purified proteins were flesh frozen in liquid nitrogen and stored at −80°C until use. Protein concentration was determined by measuring the absorbance at 280 nm using a Nanovue spectrometer and the extinction coefficient of 7450 M^-1^ cm^-1^.

To confirm the identity of htau40, protein LCMS analysis was conducted on the mass spectrometer Xevo G2-S coupled with UPLC system. The column used for the LC was Acquity UPLC Protein BEHC4 with dimensions 2.1 mm X 50 mm. The gradient for the method was: Time 0 min, Flow rate 0.2 mL/min for composition 95% A solution (Water with 0.1% Formic acid) and 5% B (Acetonitrile), Time 1.00, 95% A and 5% B, Time 5 min, 0% A and 100% B, Time 6 min 0% A and 100% B and time 7 min 95% A and 5% B. The data were processed by MassLynx software that controls LCMS analysis and runs Xevo. See additional data and figures (**Supplementary Fig. 3**) for the protein sequence and purification result of htau40.

#### MEA recordings

MEA recordings were performed using a Multi Channel Systems recording setup (Multi Channel Systems, Germany). A microscope stage-top incubator (OKOLab, Italy) was placed directly onto the MEA head stage during recording sessions and was maintained set at 37°, 5% carbon dioxide, and 20% relative humidity. On DIV 21, electrophysiological recordings were obtained. MEAs used are a mix of Indium tin oxide (ITO) MEAs devices (Multi Channel Systems MCS GmbH, 2019) and PEDOT:PSS MEAs developed in the group (Middya *et al*., 2021). First, a baseline recording 30 minutes of spontaneous activity was acquired. Subsequently, cells were treated for 2.5 hours with one of the treatment conditions, *i*.*e*., PBS (*Control*), PBS (*Stimulation-only*), 1 μM Tau (*Tau-only* and *Tau + Stimulation*). For *Control* and *Tau-only*, the cells were kept in an incubator (37°, 5% carbon dioxide, and 20% relative humidity) during the treatment period. For *Stimulation-only* and *Tau + Stimulation*, cells were kept in the MEA head stage with OKOLab setup and subjected to the stimulation protocol for the full treatment period. After treatment, another 30 minutes of spontaneous activity was recorded. For the stimulation (used for stimulated activity measurements as well as for treatment stimulation), we used three 3 monophasic pulses (amplitude −700 mV) of 200 *μ*s length and 33 ms intervals between each pulse. These stimulation pulses are then repeated every 10 seconds.

#### Spike detection and pre-processing of MEA recordings

Experimentally acquired raw MEA recordings were first filtered. As a common practice, bandpass filtering between 300 and 3000 Hz was applied. Subsequently, spike events were detected and isolated from the extracellular recording via thresholding. Thresholding was based on an estimate of the background noise σ_*m*_:

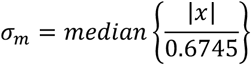

where x is the bandpass filtered signal. The threshold condition was then defined as (Quiroga, Nadasdy and Ben-Shaul, 2004):

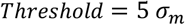

Usually, 64 sampling points were extracted for each spiking event (20 sampling points before the threshold event plus 44 sampling points after the event). The detected spike events were aligned based on the occurrence time or their minimum amplitude and further pre-processed: all spikes were min–max normalised to a scale of 0 to 1 and mapped to their gradient as it was more amenable to signal processing (Manton *et al*., 2013). Every spike recording *x*_*i*_(*t*) of the dataset *X =* {*x*_1_, *x*_2_, …, *x*_*n*_}, *x*_*n*_ ∈ *R*^*d*^ was mapped to a gradient representation ∇*x*_*i*_(*t*) as:

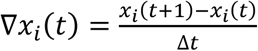

where *t* is the sampling time of each sampling point *t* ∈ (0, *d* − 1) and Δ*t* is the sampling step time (50 μs for recordings obtained at a frequency of 20 KHz).

#### Patch clamp

For patch clamp recordings, primary hippocampal cultures were prepared from postnatal day 0 or 1 (P0/P1) pups from Grik4-cre mice crossbred with Ai32(RCL-ChR2(H134R)/EYFP) (Jackson Laboratory, USA) that express ChR2-EYFP in CA3 hippocampal neurons. Recordings were carried out after 14–21 days *in vitro* (DIV) depending on the expression levels of ChR2-EYFP (DIV14–21). Coverslips were incubated and stimulated applying the following conditions: control, light stimulation, Tau (1 μM), or Tau (1 μM) + light stimulation. For the stimulation, we used three 3 monophasic pulses of 1 ms length and 33 ms intervals between each pulse. Slightly longer pulses than the electrical stimulation in MEA experiments (1 ms compared to 200 *μ*s, respectively) were chosen as required to evoke reliable responses. These stimulation pulses are then repeated every 10 seconds. Individual coverslips were transferred to an immersion-type recording chamber and perfused with artificial cerebrospinal fluid (aCSF) (126 mM NaCl, 3 mM KCl, 26.4 mM, NaH2CO3, 1.25 mM NaH2PO4, 2 mM MgSO4, 2 mM CaCl2, and 10 mM glucose, pH 7.2 and osmolarity 270–290 mOsm L−1). Patch pipettes were made from borosilicate glass capillaries (0.68 mm inner diameter, 1.2 mm outer diameter) (World Precision Instruments, UK) using a P-97 Flaming/Brown Micropipette Puller (Sutter Instrument, U.S.) with tip resistances of 4–7 MΩ. Neurons were visualised and selected using infrared differential interference contrast (DIC) microscopy using a 40x water-immersion objective. EYFP-negative neurons were identified using an U-RFL-T mercury light source (Olympus, Japan) with excitation filter 490–550 nm through the objective and selected for whole-cell patch clamp recordings in voltage clamp mode. Excitatory postsynaptic currents (EPSCs) were evoked by light stimulation (single 1 ms pulses, repeated every 20 seconds) of CA3 neurons using a DPL-473 laser controlled by a UGA-40 point laser system (3.5 - 5mW laser intensity, Rapp OptoElectronic, Germany). Data were acquired using an ITC18 interface board (Instrutech, U.S.). At least 10 EPSC traces were averaged per cell and the amplitude was analysed using Igor Pro software (WaveMetrics, U.S.).

#### Statistical analysis

GraphPad Prism 9.5.1 was used for all statistical evaluations. All datasets and comparisons have been tested for normality as well as homogeneity of variances using a Shapiro-Wilk and F-test, respectively. Accordingly, means have been compared using two-sided Student’s t-tests. For the comparison between the MEA datasets for Control and Tau condition **(Fig. 4a)**, a Welch’s t-test was applied.

## Additional Data and Figures

**Supplementary Figure 1:**
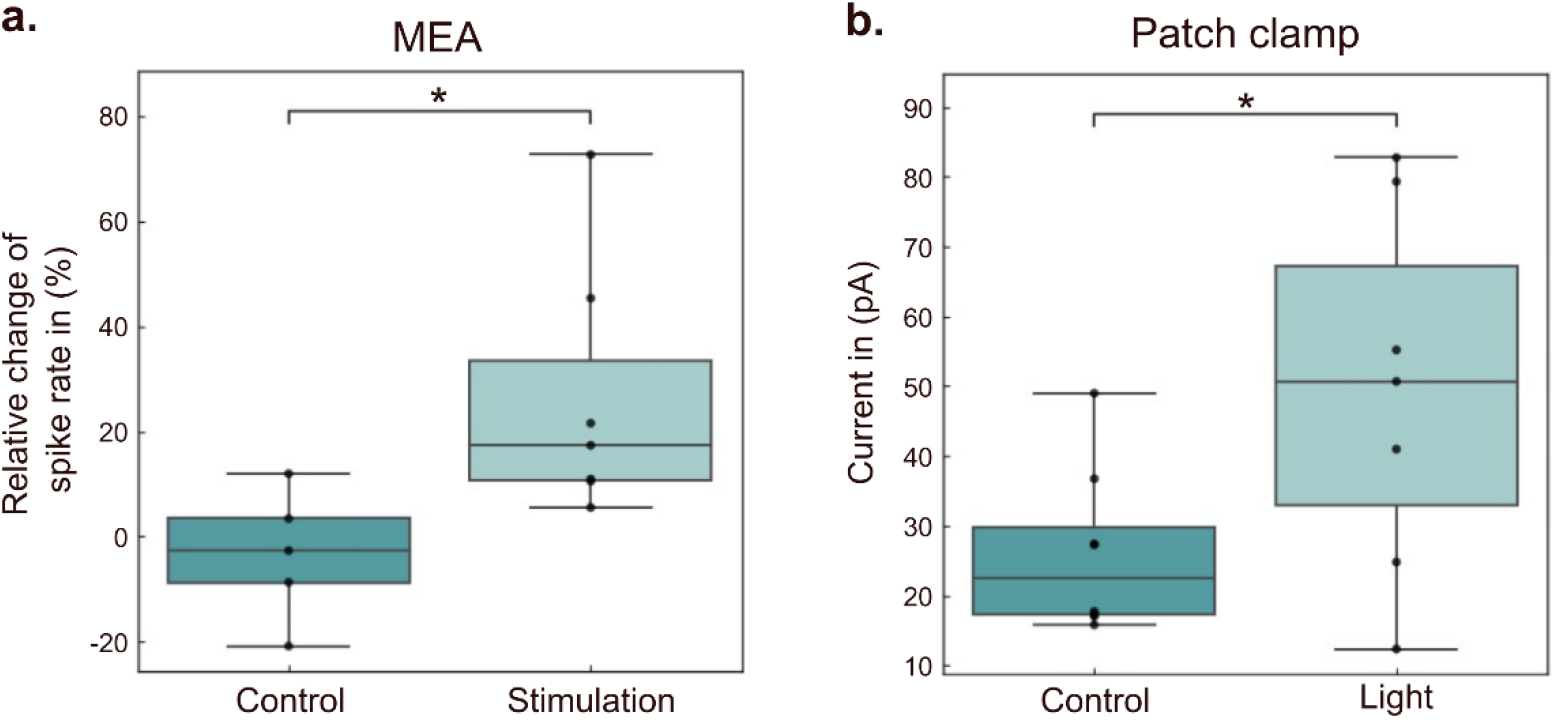
Stimulation increases activity and EPSCs in hippocampal neurons. **(a)** Relative change of spike rate measured on MEAs in the presence or absence of electrical stimulation, normalised to the pre-treatment baseline. N>=6. *P ≤ 0.05. **(b)** Patch clamp data of hippocampal neurons in the presence or absence of light stimulated activity. N>=7. *P ≤ 0.05.

**Supplementary Figure 2:**
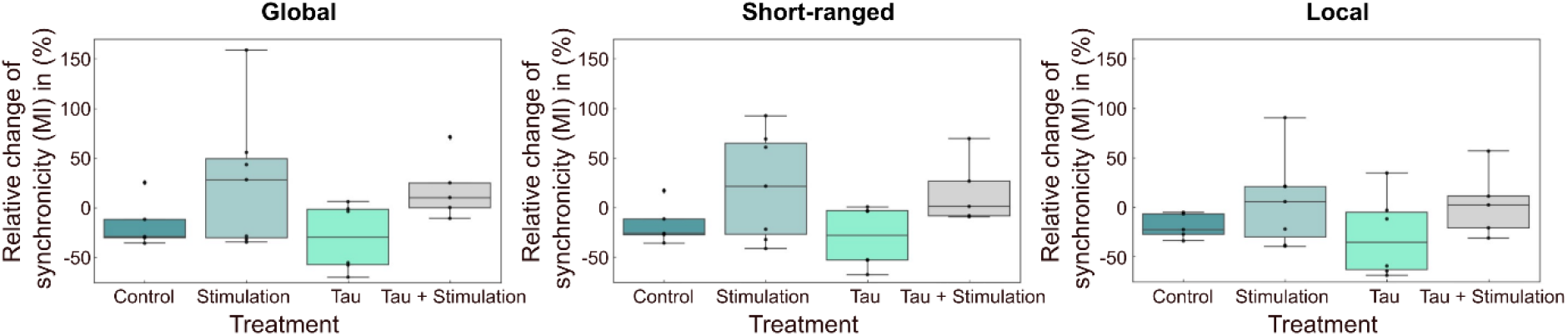
Spike Sorting enables multi-level synchronicity analysis. Shown are boxplots of the relative change in synchronicity (measured as mutual information (MI) (Gelfman *et al*., 2018)) after treatment compared to before treatment baseline. *Global* synchronicity measures the mutual information between all pairs of electrodes. *Short-range* synchronicity measures the mutual information between only neighbouring pairs of electrodes. *Local* synchronicity, enabled by spike sorting, describes mutual information between each class of detected neurons for every single electrode. Means of each electrode pair or across all electrodes (for local) are shown.

**Supplementary Figure 3:**
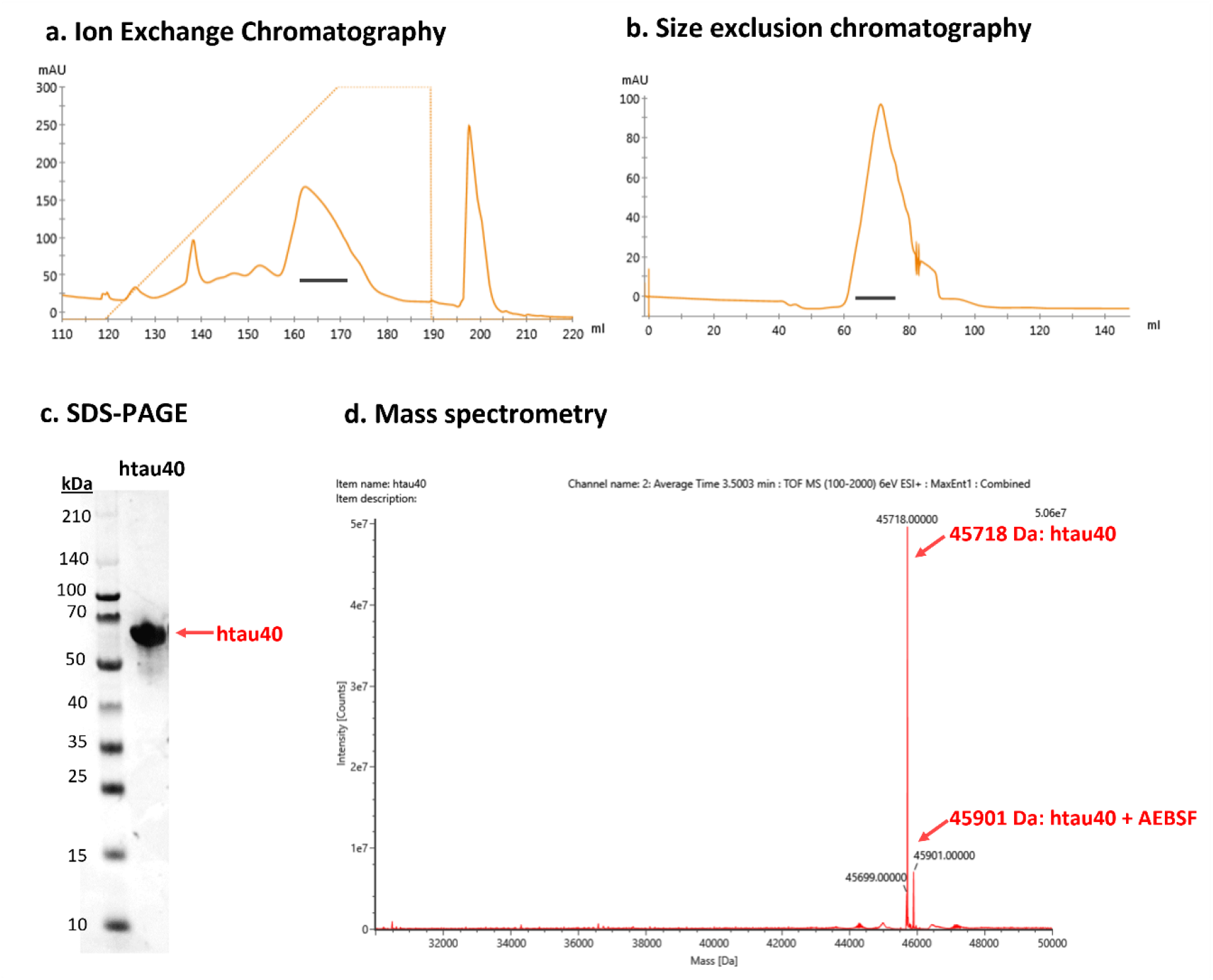
Sequential purification and analysis of htau40. **(a)** Ion-exchange chromatography using HiTrap Q HP 5 ml column with a salt gradient depicted by dashed lines for protein elution, and collected protein fractions marked by black solid lines. **(b)** Further purification via size exclusion chromatography on a Superdex 200 16/60 column with collected fraction similarly denoted by solid black lines. **(c)** Analysis of the resulting protein by SDS-PAGE to verify purity. Note that its anomalous mobility (it appears to migrate as a 60 kDa protein even though its actual mass is approximate 46 kDa) is due to its intrinsically disordered nature. **(d)** Identification and characterisation of htau40 by mass spectrometry. The 45718 Da is corresponding to the full-length htau40 without N-terminal methionine (N-terminal rule), and 45901 Da indicates the free cysteine on htau40’s interacted with AEBSF, an element of the cOmplete^TM^ protease inhibitor cocktail.

### Protein sequence htau40 (the first methionine is cleavaged due to N-dragon rule)

MAEPRQEFEVMEDHAGTYGLGDRKDQGGYTMHQDQEGDTDAGLKESPLQTPTEDGSEEPGSETSDAKSTPTAEDVTAPLVDEGAPGKQAAAQPHTEIPEGTTAEEAGIGDTPSLEDEAAGHVTQARMVSKSKDGTGSDDKKAKGADGKTKIATPRGAAPPGQKGQANATRIPAKTPPAPKTPPSSGEPPKSGDRSGYSSPGSPGTPGSRSRTPSLPTPPTREPKKVAVVRTPPKSPSSAKSRLQTAPVPMPDLKNVKSKIGSTENLKHQPGGGKVQIINKKLDLSNVQSKCGSKDNIKHVPGGGSVQIVYKPVDLSKVTSKCGSLGNIHHKPGGGQVEVKSEKLDFKDRVQSKIGSLDNITHVPGGGNKKIETHKLTFRENAKAKTDHGAEIVYKSPVVSGD TSPRHLSNVSSTGSIDMVDSPQLATLADEVSASLAKQGL

